# The how, when, and what of odor valence communication between the olfactory bulb and piriform cortex

**DOI:** 10.1101/2023.11.02.564696

**Authors:** Frans Nordén, Behzad Iravani, Martin Schaefer, Anja L. Winter, Mikael Lundqvist, Artin Arshamian, Johan N. Lundström

## Abstract

A core function of the olfactory system is to determine an odor’s valence. The central processing of odor valence is initiated in the olfactory bulb, but the neural mechanisms by which this important information is communicated to, and from, the olfactory cortex (piriform cortex) in humans are not known. To assess communication between the two nodes, we simultaneously measured odor-dependent neural activity in the olfactory bulb and piriform cortex from human participants while obtaining trial-by-trial valence ratings. We determined *when* valence information was communicated, *what* kind of information was transferred, and *how* the information was transferred (i.e., in which frequency band). Support vector machine learning on the coherence spectrum and frequency-resolved Granger causality were used to identify valence-dependent differences in functional and effective connectivity between the olfactory bulb and piriform cortex. We found that the olfactory bulb communicates odor valence to the piriform cortex in the gamma band shortly after odor onset, while the piriform cortex subsequently feeds valence-related information back to the olfactory bulb in the beta band. Decoding accuracy was better for negative than positive valence, suggesting negative valence superiority. Critically, we replicated these findings in an independent dataset using other odors across a larger perceived valence range. Combined, these results demonstrate that the olfactory bulb and piriform cortex communicate levels of odor pleasantness across multiple frequencies, at specific time-points and in a direction-dependent pattern in accordance with the two-stage model of odor processing. It also provides further evidence that odor valence should be viewed as two perceptual dimensions and not one continuous.

## INTRODUCTION

A key feature of our perceptual systems is to assign hedonic value, or valence, to stimuli to aid our decision to either approach or avoid them (Darwin, 2011; Lavender & Hommel, 2007; Öhman, 2007). In line with this, it has been hypothesized that the coding of valence is a dominant feature of the human olfactory system (Khan et al., 2007) and several areas of the olfactory system in animal models produce valence-dependent responses (Blazing & Franks, 2020; Joussain et al., 2011; Kermen et al., 2021; Secundo et al., 2014). Likewise, in humans, both the olfactory bulb (OB) (Iravani et al., 2021a) and piriform cortex (PC) (Kato et al. 2022) process representations of odor valence with inputs from the extended olfactory cortex (Rolls et al., 2003). However, how, when, and what kind of odor valence information is transferred between the nodes in the human olfactory system is not yet known.

Communication within the olfactory system occurs via the synchronization of oscillatory signals (Adrian, 1942, 1950; Freeman, 1959, 1972, 1974). Oscillatory signals between the OB and the PC in rodents have been shown to contain odor perception-related information mainly in two frequency bands, the gamma and beta band. Generally, higher frequencies, such as gamma, are often associated with processing within areas and afferent ‘bottom-up’ communication whereas lower frequencies, such as beta, are associated with efferent ‘top-down’ communication (André Moraes Bastos et al., 2015; Buschman & Miller, 2007; Frederick et al., 2016; Richter et al., 2017) In line with this are observations that odor stimulation produces coherence between OB and PC in the beta band that is mainly induced by efferent communication (Gray & Skinner, 1988; L M Kay & Freeman, 1998; Martin et al., 2006; Neville & Haberly, 2003). Although animal models have provided vast insight into the olfactory system, they are limited in their ability to directly assess the communication of odor valence between the OB and PC without dependence on some form of conditioned learning. Only indirect valence perception-dependent evidence can be obtained in animal models given their inability to verbalize their percept. However, beta activity is observed in the communication between OB and PC when an animal is subjected to aversive conditioning (Chapuis et al., 2009), which is subsequently reduced in amplitude when the centrifugal connection between OB and PC is removed, thereby suggesting that beta activity is linked to the animal’s valence perception (Martin et al. 2006).

Given the inaccessibility of the healthy human olfactory system using electrophysiological methods, little is known about how it communicates odor information. To date, only one study has assessed how the human OB and PC communicate information. Using reconstruction of the source signal based on concurrent electroencephalogram (EEG) and electrobulbogram, (EBG). Iravani and colleagues (2021) demonstrated that in response to an odor stimulus, the OB’s afferent communication to the PC is dominated by oscillations in the gamma and beta bands, whereas the PC input to the OB is dominated by theta band oscillations. These findings are consistent with computational models of the communication between the OB and PC (Chen & Padmanabhan, 2022) and with intracranial recordings from within the human PC, where oscillatory patterns induced by odor stimulation are related to theta (Jiang et al., 2017), beta, and gamma activity (Yang et al., 2022).

We recently demonstrated that the OB processes odor valence in the gamma and beta bands in a time-dependent manner with mainly early occurrence of gamma and later occurrence of beta (Iravani et al., 2021a). Because gamma oscillations in past studies have been linked to information projected from the OB (Gray & Skinner, 1988; Martin & Ravel, 2014), we hypothesized that the demonstrated gamma activity is information that flows toward the PC, whereas the beta oscillation is mainly information projected from the PC to OB. Here, we wanted to determine *the when* (time of communication), *the how* (frequencies of communication) and *the what* (pleasantness and unpleasantness) of odor valence communication between the OB and PC. To this end, we assess neural communication between the OB and PC in relation to subjective odor valence in two experiments using concurrent EBG and EEG recordings. This method allows direct, noninvasive measures from the human OB and PC (Iravani et al., 2020; Iravani, et al., 2021b). We here operationalize neural communication as coherence in neural oscillations between the two nodes (Kay et al., 2009). Specifically, we hypothesize that subjective odor valence would be communicated in early gamma activity in the direction from the OB to PC and convey primarily negative valence to facilitate a fast avoidance response. This would be followed by a later beta activity from PC to OB, which would also contain information about the pleasant percept. In Experiment 1, we determined communication of subjective odor valence between the nodes by exposing participants to two distinct classes of odors, one with a clear positive percept and one with a clear negative. In Experiment 2, we replicated findings from Experiment 1 with a new set of odors and excluded confounding contrast effects by exposing participants to the full valence range by also including odors with a neutral valence percept. Moreover, to increase decoding accuracy on the individual level, we increased the number of trials.

## MATERIALS AND METHOD

### Participants

We assessed neural responses to odors using recording of EBG and EEG signals in two separate experiments. In Experiment 1, 55 individuals participated. Due to reasons explained below, 15 participants were excluded from data analyses meaning that the final sample consisted of 40 participants (mean age 26 ± 4.5 SD; 20 women). In Experiment 2, 29 individuals participated. Due to reasons explained below, 9 participants were excluded from data analyses meaning that the final sample consisted of 20 participants (mean age 29 ± 7 SD; 7 women). In both studies, we excluded smokers and individuals with a history of head-trauma or neurological diseases. Absence of functional anosmia was determined using a screening cued odor identification test with 5 easily identifiable odors, each matched with 4 alternatives, with at least three correct answers as a cut-off. Both studies were approved by the local ethical review board (EPN: 2016/1692-31/4) and all participants signed informed consent prior to participation.

### Odor delivery and odor stimuli

In both experiments, odors were delivered for 1 second per trial using a computer controlled olfactometer with a birhinal airflow of 3 liters per minute inserted into an ongoing constant airflow of 0.5 liters per minute of clean air; a method known to eliminate potential tactile sensation of odor onset. The olfactometer has an onset time, i.e., time from trigger to 50% odor concentration at the nasal epithelium of approximately 200 ms, verified before and after each study with a photo-ionization detector (Aurora Scientific, Ontario). To enable odor presentation time-locked to nasal inhalation without obtaining attention-dependent contingent negative variation artifacts of the EEG recording, odor delivery was, unbeknown to themselves, triggered by participants’ nasal breathing cycle at the nadir of their inhalation phase, as measured by a temperature probe inserted into their nostril (sampling rate, 400 Hz; Powerlab 16/35, ADInstruments, Colorado).

Odor concentrations were determined based on results from separate pilot studies. The aim was to produce iso-intense sensations where odor laterality tests determined absence of trigeminal sensations for the presented concentrations (Hummel, 2000; Wise et al., 2012). In Experiment 1, four individual odors were presented 20 times each, rendering a total of 80 trials. These odors were Ethyl Butyrate (0.25% volume in volume dilution [v/v], Sigma Aldrich, CAS 105-54-4), Diethyl Disulfide (0.25%, Sigma Aldrich, CAS 110-81-6), Carvone (50%, Merck, CAS 6485-40-1), and Fish odor (50%, odor mixture from Symrise Inc). In Experiment 2, six individual odorants were presented 30 times each, rendering a total of 180 trials. These odors were Linalool (0.14% v/v, Sigma Aldrich, CAS 78-70-6), Ethyl Butyrate (0.25% v/v, Sigma Aldrich, CAS 105-54-4), 2-Phenyl-Ethanol (0.1% v/v, Sigma Aldrich, CAS 60-12-8), 1-Octen-3-OL (0.2%, Sigma Aldrich, CAS 3391-86-4), Octanoic Acid (1%, Sigma Aldrich, CAS 124-07-2), and Diethyl Disulfide (0.25%, Sigma Aldrich, CAS 110-81-6). All odors in both experiments were diluted in neat diethyl phthalate (99.5% pure, Sigma Aldrich, CAS 84-66-2).

Although consistency across individuals do exist in respect of odor valence, assigning valence based on odorant for a group is inherently problematic given the large individual variation originating from past experiences and associations to the odor in question. To circumvent this problem, in all our analyses, valence grouping was based on rated subjective valence for each trial without taking odorant into account. In addition, for each participant, trials were divided into three equal categories based on rated valence, with the two extreme categories serving as pleasant (top ⅓ rated trials) and unpleasant (bottom ⅓ rated trials) odor sensations. This approach assured a clear separation between categories based on subjective experience rather than odorant. Moreover, the separation of pleasant and unpleasant trials aligns with the notion that there is not a continuous odor valence dimension. Instead, it has been proposed that odor valence acts along two dimensions; one pleasant and one unpleasant, which serve the role of affect classification rather than a fine-tune valence scale (Rouby & Bensafi, 2002; Russell, 1980). This hypothesis finds support in our past work where odors seem to be processed within broad valence categories rather than along a continuum (Iravani et al., 2021a). In both experiments, there was a clear separation in rated valence perception between categories; both considering within the individual and group effects (Figure 2; Experiment 1: two-tail *t*-test. *t*(20) = 9.6, *p* < .0001, CI = [39.99, 50,99]; Experiment 2: *t*(40) = 17.2, *p* < .0001, CI = [28.19, 44.08]).

**Figure 1.**
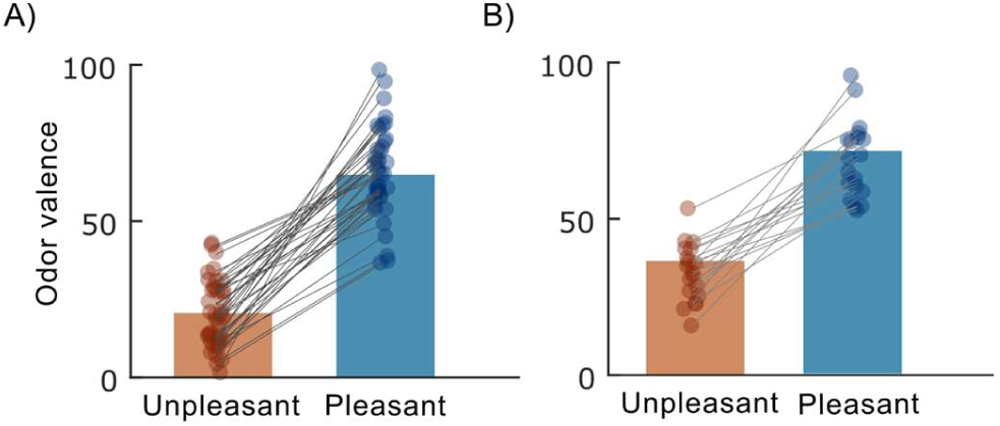
Mean rated valence for odor stimuli. **A**) Bars indicate group averaged odor valence ratings for included trials in Experiment 1, SD = 11 vs 14 for the unpleasant and pleasant classification. **B**) Bars indicate group averaged odor valence ratings for included trials in Experiment 2. SD= 9 vs 12 for the unpleasant and pleasant classification. In both panels, participant’s ratings are marked with filled circles for both odor categories (i.e., unpleasant and pleasant), which are connected with grey lines. Overlapping individual ratings are slightly shifted sideways for increased visibility.

**Figure 2:**
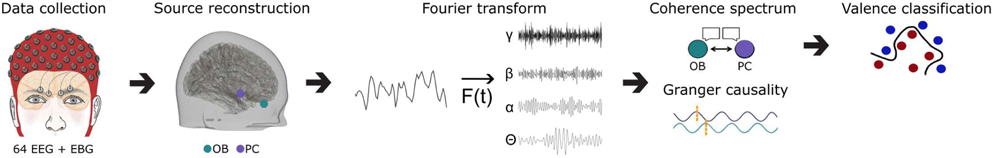
Experimental setup and analysis pipeline. EEG and EBG recordings were collected from healthy participants. Valence was rated after each trial and the data was source reconstructed to extract the signal from the OB and PC. The phase and amplitude were extracted from the source reconstructed time course via the Fourier transform and used to calculate the Coherence spectrum and frequency resolved Granger causality between the OB and PC. We furthermore applied a Support Vector Machine classifier on the Coherence spectrum to distinguish coherence related to valence.

### Testing procedure

Participants were tested in a sound-attenuated and well-ventilated chamber, specifically designed for olfactory-EEG testing. To prevent auditory cues that might alert participants to the delivery of an odor, participants wore headphones that continuously played low volume white noise. Experiment 1 consisted of four testing blocks and Experiment 2 three. Each testing block lasted 15 minutes and was separated by 5-minute breaks to reduce odor adaptation and habituation and to limit participant fatigue. Event timing and odor triggering were implemented using E-prime 2 (Psychology Software Tools, Pennsylvania). Odors were presented in a randomized order with balanced odor presentation between testing blocks. After each odor presentation, participants rated how pleasant and intense they perceived the odor on a visual analogue scale, ranging from 0 (very unpleasant/very weak) to 100 (very pleasant/very strong).

### EEG and EBG recording settings

Neural responses to the odor stimuli were acquired at 512 Hz using 64 scalp and 4 EBG active electrodes with ActiveTwo system (Bio-Semi, Amsterdam, The Netherlands). Prior to the experiment, the offset of all electrodes was manually checked, and any electrodes with an offset greater than 40 μV were manually adjusted. Stereotactic coordinates of all electrodes were acquired using an optical neuronavigation system (Brainsight, Rouge Research, Montreal, Canada). The digitalization protocol involved localizing fiducial landmarks such as the nasion and left/right preauricular points, as well as the central point of each electrode. These landmarks were then used to co-register each electrode to the standard MNI space. The digitalized electrode positions were later used in the eLORETA algorithm to enable the localization of signal sources (see below).

### Preprocessing

The data was epoched to 2000 ms long segments, from 500 ms pre-stimulus to 1500 ms post-stimulus and re-referenced to the average activity of all electrodes. Power line interference (50 Hz) was filtered out with a discrete Fourier transform filter. Eye blinks were removed with Independent Component Analysis with the InfoMax algorithm (Zhaojun Xue et al., 2006) and large muscle movements were detected by extracting z-scored Hilbert transform amplitude values. Trials with a z-score above 7 were removed from further analysis. Furthermore, participants who had more than half of the trials eliminated in one or more categories during the preprocessing were left out from further analysis. The final sample for Experiment 1 comprised of 40 participants with an average of 59 (±19) remaining trials, and the final sample for Experiment 2, comprised of 20 participants with an average of 145 (±47) remaining trials.

### Signal processing and data reduction

#### Source time-course reconstruction

To reconstruct the source-time course for the OB and PC dipoles, we first used the digitized electrodes to co-register the participants’ head to default MNI space using a six-parameter affine transformation. The MNI-152 template was subsequently used to calculate the forward model through the Finite Element Method, following the description in Fuchs et al. (2007). The T1 scan was segmented into five materials, CSF, gray matter, white matter, scalp, and skull with conductance’s [0.43, 0.01, 1.79, 0.33, .14] (Hallez et al., 2007). A Freesurfer generated cortex model, built on icosahedrons with resolution 7, was used to create the source model. This source model was used to attain the source activity through solving the inverse problem with eLORETA with a regularization parameter set to 10 %. Finally, singular value decomposition was used to project the source activity to the principal axis. The analysis was later constrained to four ROIs where the dipoles correspond to left *(x−4, y+40, z−30)* and right OB *(x+4,y+40,z−30)*, determined based on T2 weighted images, as well as left *(x -22, y+0, z -14)* and right PC *(x+22, y+2, z -12)* (Seubert et al., 2013). The source reconstruction was performed with Fieldtrip toolbox 2022 within Matlab 2022a (Oostenveld et al., 2011).

#### Source connectivity

When investigating the connectivity between neuronal populations, two approaches can be taken: functional or effective connectivity. Functional connectivity determines the relationship in amplitude or phase between two neuronal populations (André M Bastos & Schoffelen, 2015; Biswal et al., 1995), while effective connectivity determines the predictive relationship between the two (Eldawlatly & Oweiss, 2010). To assess the connectivity between OB and PC, both functional and effective connectivity was used to form a complete picture containing both phase relationship and directionality.

For the functional connectivity, the source reconstructed time course is transformed into Fourier space where the coherence spectrogram is used to evaluate where linear information transfer occurs between the OB and PC. To evaluate the effective connectivity between the OB and PC, an approach based on spectral Granger causality was used. Granger causality assesses whether the future of a time series (X) can be predicted by past values of X alone or if it is more accurately predicted by past values in the alternative time series (Y) (Granger 1969). Spectral Granger causality builds upon the same concept, but the assessment is here performed in Fourier space, where a transfer function is calculated for the source-reconstructed time course. Due to our interest in the frequency domain, we used spectral Granger causality to evaluate the directionality of communication between the OB and PC.

#### Functional connectivity in frequency and time

The OB and PC coherence spectral density was calculated between 4-100 Hz with a multi-tapering convolutional method with two tapers of Discrete Prolate Spheroidal Sequences (DPSS), combined with a flexible time-window. This enabled high frequency resolution in the lower frequencies, and at the same time, the possibility to represent at minimum two cycles per time-frequency window. The applied frequency smoothing was set to 80 % of the target frequency to enable good frequency resolution in all bands while maintaining a time resolution that considers the variability in timing between trials.

#### Effective connectivity in the frequency domain

Transformation to Fourier space for the source reconstructed time course was estimated within the frequency range 4-100 Hz with a multi-tapered fast Fourier transform. The step was set to 1 Hz with the smoothing parameter to 5 Hz with 7 DPSS tapers and applied to the whole stimulus time period, i.e., 1 second. High frequency accuracy could therefore be achieved by choosing a low smoothing parameter. The Spectral Granger causality was averaged over hemispheres to increase statistical power. Statistical significance was determined on a group level by applying a two-tailed Student’s *t*-test.

#### Support Vector Machine learning

To determine whether odor valence information could be assessed from the communication between the OB and PC, a Support Vector Machine (SVM) classifier was applied to the OB-PC coherence spectrogram. The entire coherence spectrogram was searched iteratively to determine whether any region of the CS could be used to predict odor valence from the functional communication between OB and PC using SVMs. Quadratic areas of 121 samples around each point in the coherence spectrogram were chosen as the assessed area and binned together to create the feature space. The whole cross-spectrogram was subsequently evaluated in a searchlight manner. Bins with less than 10 neighbors were excluded from further analysis. A leave-one-out scheme was applied on the data where each participant was left out once per quadratic area. The mean accuracy on the group level was compared with the chance level for two classes through a non-parametric statistics 5000 permutations Monte-Carlo test. Confidence intervals provided as 95% confidence range.

## RESULTS

### Gamma and beta activity communicates odor valence between OB and PC

We first wanted to determine whether information related to odor valence could be extracted from the communication between OB and PC around the time of odor stimulation. To do this, we applied an SVM on the coherence spectrogram to decode valence from the functional connectivity between the OB and PC. The SVM was applied to a binary classification problem with one class containing the designated unpleasant trials and the other class containing the designated pleasant trials. In Experiment 1 (Figure 3A), we were able to extract information about valence above chance levels in a broad band gamma frequency (∼50-100Hz) at around 150 ms, 400 ms, and 600-700 ms from odor onset. In addition, we were able to extract significant information in the beta frequency, but in a more diffused fashion with high decoding values mostly at later time points, around 600 ms and 850 ms after odor onset.

**Figure 3:**
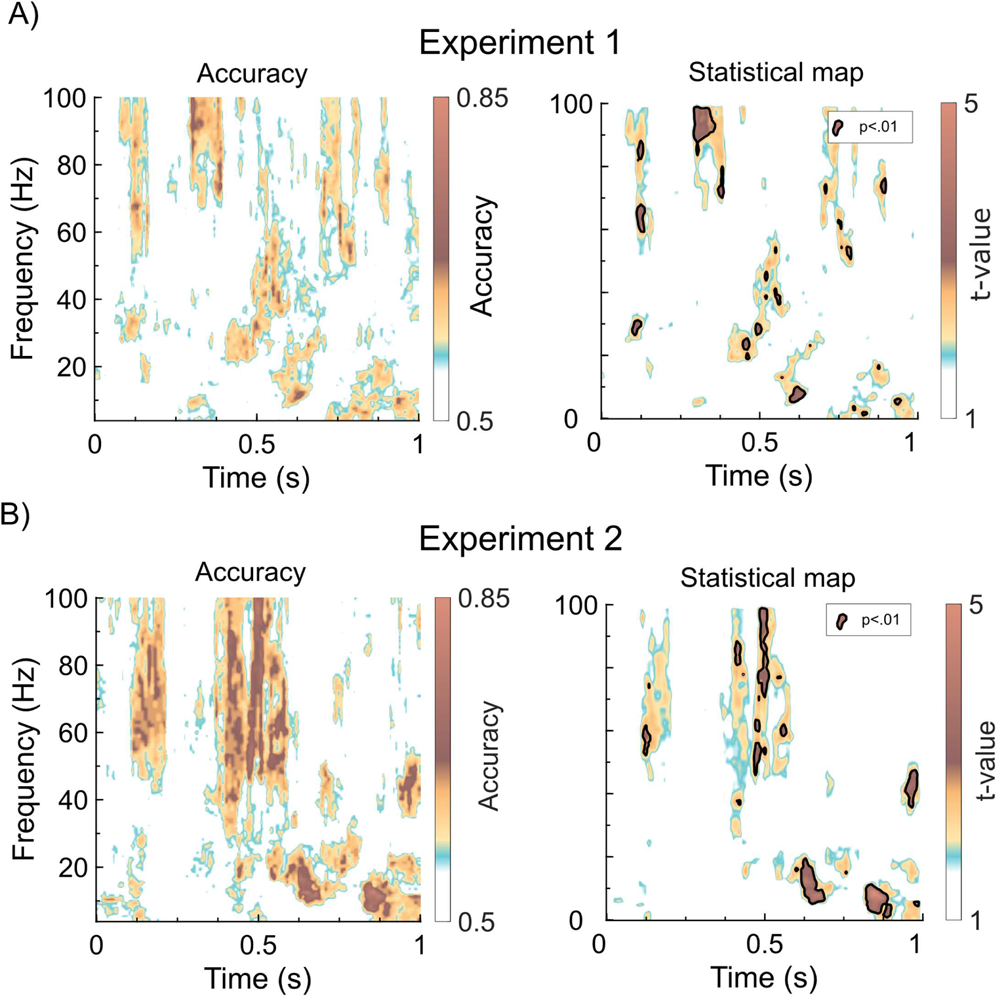
Consistent decoding of odor valence in gamma and beta connections between OB and PC across both studies. **A)** Left panel: Classification accuracy in the coherence spectrum between OB and PC for Experiment 1. Right panel: *t*-value map of the classification accuracy between OB and PC, where significant (*p* < .01) clusters of decoding accuracy above chance level are marked with black borders. **B)** Corresponding plots for Experiment 2.

Having demonstrated that valence-related information could be extracted from the coherence spectrum, we assessed the decoding accuracy. We found that in the early gamma area, around 68 Hz, we achieved a 67% decoding accuracy, *t*(40) = 3.45, *p* = .0013, *CI-range* = .001. In the beta activity, around 600 ms at 12 Hz, the highest classification accuracy was 70%, *t*(40) = 3.5, *p* = .0011, *CI-range* = .0009. Moreover, in the late beta region around 900 ms in, we had an accuracy of 70% at 12 Hz, *t*(40) = 3.1, *p* = .0035, *CI-range* = 0.0016.

The results from Experiment 1 were largely replicated in the subsequent Experiment 2 (Figure 3B), where the increased decoding power on the individual level yielded similar but more coherent results. Once again, significant decoding accuracy was found in the gamma band (∼40-100 Hz) at a time around 150 ms, 400-600 ms, and a small burst around 1000 ms from odor onset. In the beta band, two clear areas of significant decoding appear around 600 ms and 850 ms from odor onset. Assessing decoding accuracy, we found that in the early gamma region, we achieved a maximum classification accuracy of 68% at a frequency around 63 Hz, *t*(20) = 3.1, *p* = .0056, *CI-range* = .002. For the beta activity at 20 Hz around 600 ms into the trial, we had a decoding accuracy of 75%, *t*(20) = 4.5, *p* = .00022, *CI-range* = .000041. The highest decoding accuracy was found in the late beta activity around 14 Hz at 850 ms, with an accuracy of 85%, *t*(20) = 5.1, *p* < .000055, *CI-range* = .00002.

Interestingly, the later region we identified in both experiments consisted of a broad band-like activity in the theta, alpha, and beta band. Significant activity in the theta range was found in Experiment 1 at 6 Hz at 850 ms, *t*(40) = 2.9, *p* = .0060, *CI-range* = .002, and at 980 ms, *t*(40) = 2.4, *p* = .021, *CI-range* = .004. This activity happened around the peak of inhalation which had a group mean = 910 ms and SD = 50 ms. In Experiment 2, we found similarly significant theta activity at 6 Hz around 900 ms, *t*(20) = 3.2, *p* = 0.0045, *CI-range* = .002, and also later at 960 ms, *t*(20) = 2.5, *p* =.021, *CI-range* = .004. Also here, the activity occurred around the peak of inhalation (mean = 900 ms, SD = 70 ms). Given the link between theta activity and both valence perception as well as breathing, we then assessed potential behavioral differences in breathing parameters between valence condition to explore a potential breathing-related confound. However, no significant differences in breathing volume between odor valence groupings were found in either Experiment 1 or 2, with a mean area under the curve for unpleasant odors being 145 (arbitrary units, SD = 90) and for pleasant odors being 142 (arbitrary units, SD = 134), *t*(40) = 0.22, *p* = .82, *CI* = [-28.3, 35.3] in Experiment 1 and in Experiment 2, unpleasant 143 (arbitrary units, SD = 97) and pleasant 133 (arbitrary units, SD = 63), *t*(20) = 0.99, *p* = .34, *CI* = [-12.0, 33.2].

We then wanted to determine how consistent the valence classification results were across individuals to understand potential individual differences. The significant regions that matched in the two experiments were the early gamma region and the two later beta regions (Figure 4A). We selected these regions as our regions of interest (ROIs) and from these, extracted the mean classification accuracy for each participant. In the ROI located within the gamma frequency around 100 ms after odor onset, we found a clear bimodal distribution with a large proportion of our classifications centered around chance, with results being very consistent between Experiments (Figure 4Bi). To disentangle our classification accuracy, we produced a confusion matrix for the same area where it became clear that we can classify unpleasant odors with higher accuracy compared to the pleasant odors, (e.g., Experiment 2: 0.74 compared to 0.57; Figure 4Ci). For the two ROIs located in the beta band (Figure 4B ii-iii), the distributions were again consistent across experiments. However, for the beta ROI located around 600 ms from odor onset, overall decoding accuracy was clearly higher in Experiment 2, whereas the distribution for Experiment 1 demonstrated a bimodal tendency. That said, we could classify both valence classes with the unpleasant class showing higher accuracy than the pleasant class with 0.8 compared to 0.63 in Experiment 2 (Figure 4Cii). Classification in the late beta was more balanced, but still in favor of the unpleasant class with 0.77 against 0.65 (Figure 4Cii).

**Figure 4:**
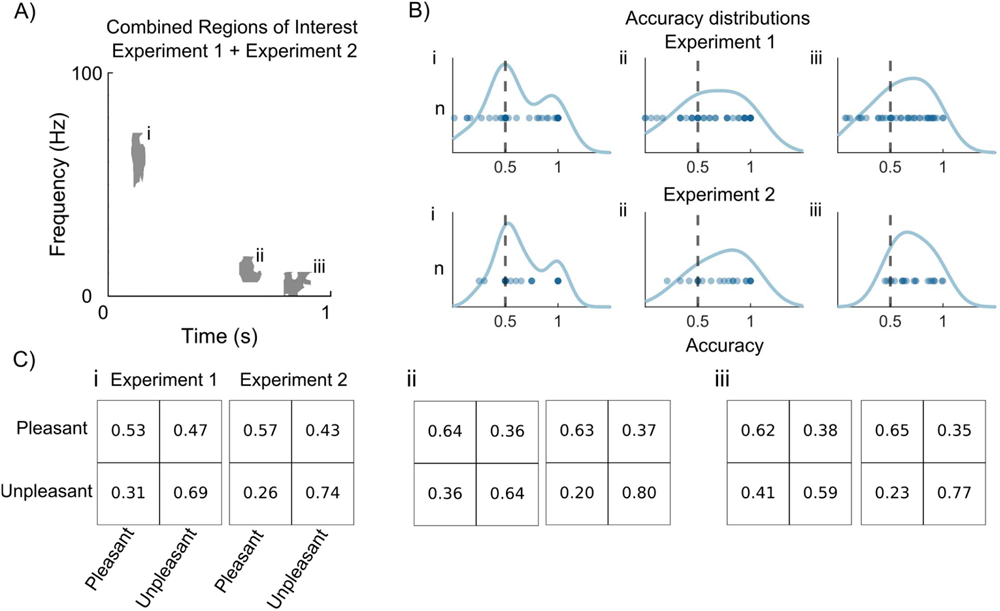
Valence separation within regions of interest. **A**) The three regions of interest, Experiment 1 overlayed Experiment 2. **B**) Distributions of classification accuracy for each participant in the three regions of interest for Experiment 1 and Experiment 2 where the dots represent participants **C**) Confusion matrices for Experiment 1 and 2 for the three regions of interest.

**Figure 5:**
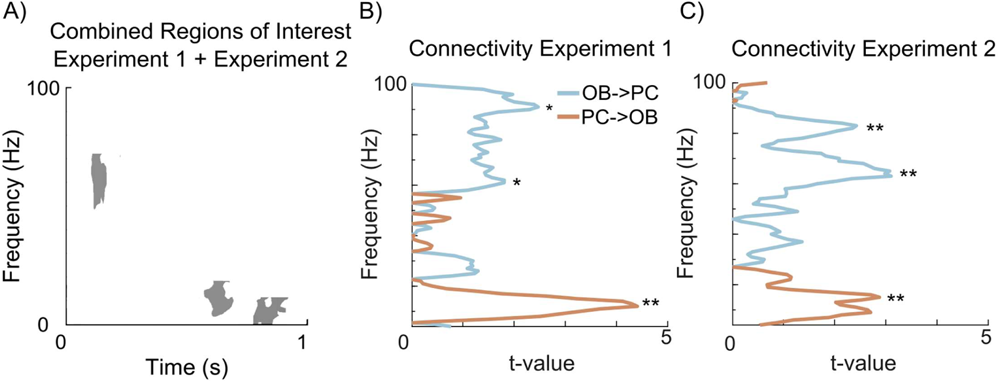
Directional communication of odor valence. **A**) The three areas of interest that were found significant and replicated in the classification analysis. **B**) Frequency resolved Granger Causality from the OB to PC (blue) and from PC to OB (orange) for Experiment 1 where we find significant clusters of information transfer *p* < .05 in gamma (OB to PC) and beta (PC to OB). * Represents *p <*.*05* and ** *p <* .*01*. **C**) Corresponding plot for Experiment 2.

Odor intensity is often considered a confounding factor in studies of odor valence. Although we aimed for intensity matched odor in our pilot studies, to assess the potential for intensity related confounds, we conducted the same analysis as described above, but classified the trials based on perceived intensity instead of valence. The results of this analysis can be found in the Supplementary material. Critically, none of the time-frequency areas identified as responding to intensity were reproduced between Experiment 1 and 2 and none of the areas identified as linked to intensity overlapped with areas linked to odor valence in either experiment.

### Effective connectivity is reciprocal between OB and PC

We next wanted to explore the potential directionality of the signals. To this end, we applied spectrally resolved Granger causality to evaluate the effective connectivity between the OB and PC in a signal-directional manner. To determine if the two nodes communicate in different bands, the frequency resolved Granger causality for the unpleasant and pleasant trials were contrasted against each other. In Experiment 1, significant information transfer was found in the direction from the OB to PC in the gamma band with peaks around 60 Hz, *t*(40) = 2.1, *p* = .0042, CI = [0.008, 0.4477], and 90 Hz, *t*(40) = 2.7, *p* < .0088, CI = [0.0698, 0.4709]. In the opposite direction, from PC to OB, we instead found significant information transfer in the beta band with a peak at 13 Hz, *t*(40) = 4.99, *p* = .0000035, CI = [0.5348, 1.2446]. This indicates that the early gamma region is related to information transferred from the OB to PC and that the late beta regions are related to information transferred from PC to OB.

For experiment 2 we found comparable results where there was a significant information transfer from the OB to PC in the gamma band with peaks at 64 Hz, *t*(20) = 3.9, *p* = .0004, *CI* = [0.2472, 0.7878], and 82 Hz, *t*(20) = 2.7, *p* = .0092, *CI* = [0.0855, 0.5681]. We also observed similar beta communication as in Experiment 1, from PC to OB, with the information transfer in the beta band, peaking at 15 Hz, *t*(20) = 3.1, *p* = .0034, *CI* = [-1.2319, -0.26219].

## DISCUSSION

Our aim was to determine *how, when*, and *what* kind of odor valence information is communicated between the OB and PC, two critical nodes of the olfactory system. Our results suggest that odor valence is reciprocally communicated between the OB and PC in the gamma and beta bands. Moreover, our results suggest that the OB communicates valence-related information to the PC in the gamma band around 150 ms after odor onset, and that the PC projects back at two timepoints in the beta band, 700 ms and 850 ms after odor stimulus presentation. Our results also suggest that the early gamma region primarily communicates negative odor valence while the later beta regions contain a richer representation of the odor valence percept. These findings demonstrate that the OB and PC communicate odor valence information across multiple frequencies at specific time-points and in a direction-dependent manner.

Our results are consistent with the general model of bottom-up/top-down modulation found in the visual and higher order cognitive systems. In this general model, higher oscillatory frequencies, such as gamma, are associated with afferent ‘bottom-up’ communication, while lower frequencies, such as beta, are associated with efferent ‘top-down’ communication. Our analysis suggests that the information transfer from OB to PC is dominated by gamma activity, while the transfer from PC to OB is dominated by beta activity. This result aligns with the identified areas in the time-frequency coherence spectra between the OB and PC found to be associated with valence processing in the two experiments.

In a previous study focusing on valence processing within the OB, we identified a period of late gamma processing in the OB that related to subjective odor valence (Iravani et al., 2021a). In the present two experiments, we identified valence-dependent activity in the beta band that overlaps in time with the aforementioned late gamma processing in the OB. Given that the final odor valence percept is highly contextdependent (Djordjevic et al., 2008) and that we find late beta activity in the direction from PC to OB, we hypothesis that information in the beta band updates the prior information in the OB with known contextual and memory-related information regarding the perceived odor, which in turn influences late gamma processing. Such updated prior information would then also modulate OB processing of any subsequent odor and form reciprocal communication within the olfactory system based on top-down context information from higher order brain regions.

Our findings of early gamma and late beta processing align with the two-stage model of odor processing (Frederick et al., 2016) which stipulates that the OB, when receiving odor information, initially processes the odor by a fast gamma response and then a later and slower beta response. Within this framework, the early gamma response contains basic information about the odor that is meant to facilitate fast discrimination and potential avoidance, while the beta activity is thought to contain a much richer representation of the odor by incorporating top-down information (Frederick et al., 2016; Iravani, et al., 2021a). We find evidence for this theory both through the timing of our areas (early gamma, later beta) and through the increase of decoding accuracy over time between the two frequency bands. The difference in classification accuracy between the early gamma and later beta activity is 17%, which is consistent with previous data from the human PC showing that the possibility to classify perceived valence increases over time (Kato et al., 2022). This difference was mainly due to early gamma predominantly containing information about negative valence, whereas later beta demonstrated good classification for both negative and positive valence. Indeed, the need for fast discrimination and action is more important for unpleasant than pleasant odors due to the ecologically higher importance of avoiding danger than approaching reward.

Recent work indicates a separation between pleasant and unpleasant odors in their neural processing (Bensafi et al., 2002; Rouby & Bensafi, 2002). Our own past work of valence processing within the OB mainly finds activation related to unpleasant odors (Iravani et al., 2021a) and other recent studies point towards the Olfactory Tubercle, a region down-stream to the OB that is highly connected to the striatum (Xiong & Wesson, 2016), as the processing point for pleasant odors in both animals (Gadziola et al., 2015; Wesson & Wilson, 2011) and humans (Midroit et al., 2021). Another region involved in odor processing, the Orbitofrontal Cortex, has during intracranial stimulation, only generated pleasant odor perception among the participants (Bérard et al., 2021). In line with the notion that positive odor valence mainly originates from outside the OB, we obtained higher decoding accuracy for pleasant odors in the late beta activity. Given that beta in our data originated from PC, our results suggest that a key driver of pleasant odor valence processing is information originating from upstream cerebral areas and not predominantly linked to receptor activity based on the chemical composition. However, it should be noted that recent data from animal models suggesting that odors associated with reward produce wide-spread inhibition of mitral cells in the OB (Lindeman et al., 2023). Also, optogenetically stimulating the posterior portion of the rodent OB is rewarding to the animal (Midroit et al., 2021). Whether methodological differences account for this discrepancy or whether there are inherent differences in how the human and rodent olfactory systems are organized remains to be determined. That said, we find that communication in the beta band is linked to valence processing, a finding that corresponds to animal experiments using aversive conditioning (Chapuis et al., 2009). Although direct comparisons and conclusions are not possible based on these experiments, the similarity in results both suggest that potential discrepancies are due to methodological differences and that the use of aversive conditioning as a tool to modulate valence is, at least in part, a good model for human valence perception.

Individual variation, followed closely by the molecular structure of the odorant, is the main factor explaining the difference in odor valence perception (Arshamian et al., 2022). Indeed, individual variation in odor valence perception is large even within families (Logue et al., 1988). Despite trying to limit individual variation by using the subjective valence percept of each individual and trial, we still see a great deal of individual variation in how well we can classify valence (Fig. 4b). However, it is worth noting that the final odor percept, i.e., the percept that is mainly formed by the top-down beta activity and reflected in the two later beta regions, was better classified than the early gamma region, which, as we have previously demonstrated, mainly codes avoidance information (Iravani, et al., 2021a). This indicates that our approach of using individual ratings instead of predetermined valence categories was useful. However, it is important to clarify that this approach does not enable us to control the subjective experiences themselves. For instance, two participants classifying an odor as negative may have entirely different subjective experiences.

In both experiments, we found significant valence-related activity in the theta band around 900 ms after odor onset. Theta is a prevalent frequency in the olfactory system and often related to respiration, with the peak in theta occurring at peak inhalation (Jiang et al., 2017). Due to this, theta activity in the PC and OB is generally considered to hold information about breathing and sniff behavior. In our data, we can distinguish between pleasant and unpleasant odor percepts in theta band coherence at the peak of inhalation. However, we did not find any difference in sniff-related parameters between the two experiments. The absence of difference in the sniff of the actual odor and the late time points of the theta activities suggests that the theta activity in our two experiments contains information regarding valence-related regulation of the next sniff. Our experimental design unfortunately prohibits us to assess potential linkage between OB-PC theta communication, sniff behavior, and odor valence in subsequent sniffs of odors. Given the tight coupling between sniff behavior and odor perception, future studies should aim to design their experiments to facilitate this comparison.

Given the inherent ethical problems of performing invasive brain recordings on healthy human subjects, we collected EEG and EBG data that we then transformed into source space. Thereby, we assess an indirect signal with weaker signal-to-noise-ratio (SNR) than intracranial measures directly from the region in question. That said, the Finite Element Model-based methods of EEG source reconstruction have been demonstrated to reconstruct subcortical sources with a reliable SNR (Piastra et al., 2021) and past studies have demonstrated that our methods allow for extraction of reliable and reproducible neural signals from both the OB (Iravani et al., 2020) and PC (Iravani, et al., 2021b) which aligns with results obtained using intracranial measures (Yang et al., 2022). The robustness of our findings is also demonstrated by the ability to replicate them in independent experiments.

Odor intensity and valence are inherently linked, with varying degrees of coupling. In our experiments, we demonstrated that intensity coding did not overlap in time and frequency band with our valence-related results. Moreover, our experiments were designed to minimize the impact of odor intensity perception, which means that we will have lower power for intensity compared to valence classification. Therefore, our findings are unlikely to be explained by intensity differences between pleasant and unpleasant odors.

In conclusion, odor valence is communicated between the olfactory bulb (OB) and piriform cortex (PC) in two principal frequency bands: gamma and beta. This communication is directional, with bottom-up information communicated in gamma and top-down information communicated in beta. Our results further suggest that odor valence is communicated between the OB and PC during two time periods. First, odor valence is communicated from the OB to the PC in gamma, starting 150 ms after odor onset. Then, the PC communicates valence-related information back to the OB in beta at two timepoints: 700 ms and 850 ms after odor onset. Finally, our results indicate that primarily the unpleasant percept is communicated in the early gamma region, while the later regions contain a richer and more balanced valence percept. These results support the two-stage model of odor processing and also provide further evidence that odor valence should be viewed as two perceptual dimensions and not one continuous.

## Supporting information

Supplementary analysis 1

## Data availability

The collected data and the code used to analyze it is available through OSF via https://osf.io/fqw2m/?view_only=4a3ca5b9c9894f91bd2165f67ffd49fa

## Acknowledgement

Funding provided by the Knut and Alice Wallenberg Foundation (KAW 2018.0152) and the Swedish Research Council (2021-06527), awarded to JNL. Data acquisition supported by a grant to the Stockholm University Brain Imaging Centre (SU FV-5.1.2-1035-15).

