## Supplementary analysis 1 for "The how, when, and what of odor valence communication between the olfactory bulb and piriform cortex"

### Supplementary material

*Intensity as a control*

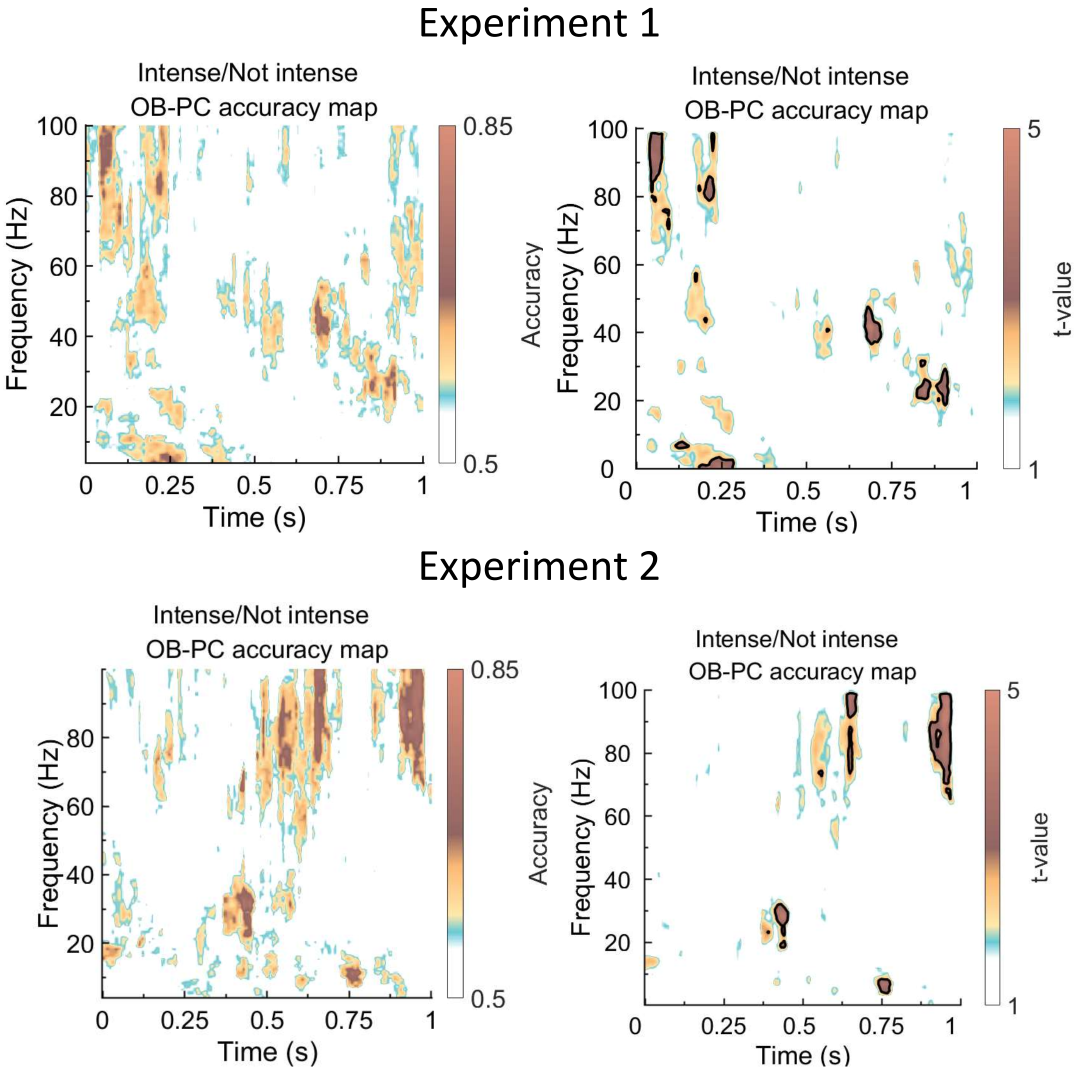


**Figure S1:** Accuracy maps when classifying intensity. No significant areas were replicated over the two studies and no overlap compared with the pleasantness maps was found. We used the same setup with the Support Vector Machine on the two classes of high intensity odor and low intensity odor. In the accuracy map seen in Figure 3 we can clearly see that the results found with regards to valence are not dependent on the intensity. While there is significant activity with regards to intensity it is not in the same regions as the one found with regards to valence.
